# Evolutionarily informed deep learning methods: Predicting transcript abundance from DNA sequence

**DOI:** 10.1101/372367

**Authors:** Jacob D. Washburn, Maria Katherine Mejia-Guerra, Guillaume Ramstein, Karl A. Kremling, Ravi Valluru, Edward S. Buckler, Hai Wang

## Abstract

Deep learning methodologies have revolutionized prediction in many fields, and show potential to do the same in molecular biology and genetics. However, applying these methods in their current forms ignores evolutionary dependencies within biological systems and can result in false positives and spurious conclusions. We developed two novel approaches that account for evolutionary relatedness in machine learning models: 1) gene-family guided splitting, and 2) ortholog contrasts. The first approach accounts for evolution by constraining the models training and testing sets to include different gene families. The second, uses evolutionarily informed comparisons between orthologous genes to both control for and leverage evolutionary divergence during the training process. The two approaches were explored and validated within the context of mRNA expression level prediction, and have prediction auROC values ranging from 0.72 to 0.94. Model weight inspections showed biologically interpretable patterns, resulting in the novel hypothesis that the 3’ UTR is more important for fine tuning mRNA abundance levels while the 5’ UTR is more important for large scale changes.

## INTRODUCTION

Machine and Deep Learning approaches such as Convolutional Neural Networks (CNNs) are largely responsible for a recent paradigm shift in image and natural language processing. They are some of the fundamental enablers of modern artificial intelligence advances such facial, recognition, speech recognition, and self-driving vehicles. The same Deep Learning approaches are beginning to be applied to molecular biology, genetics, agriculture, and medicine ^1–4^, but evolutionary relationships make properly training and testing biologically useful models much more challenging than the image or text classification problems mentioned above.

For example, if one wishes to predict mRNA levels from DNA promoter regions (as we do here), the standard approach from image recognition problems would be to randomly split genes into training and testing sets ^5^. However, such a split will likely lead to dependencies between the sets because of shared evolutionary histories between genes (i.e., gene family relatedness, gene duplications, etc.), and may cause model overfitting and false positive spurious conclusions. Models trained without properly accounting for the biology of evolution (and perhaps other biological and technical factors specific to the modeling scenario) will likely perform poorly when challenged with new data, and they will have limited, if any, utility for biological discovery. This is because they have memorized both the neutral and functional evolutionary history, rather than only learning the functional elements. Such erroneous models would lead researchers and other users to make incorrect conclusions.

With these challenges in mind, we developed two CNN architectures for predicting mRNA expression levels from DNA promoter and/or terminator regions (as defined in the online methods section). These include models that predict: 1) if a given gene is highly or lowly expressed, and 2) which of two compared gene orthologs has higher mRNA abundance. The architectures are built around two methods we developed for properly structuring the model training and testing process to avoid the issues of training set contamination by evolutionary relatedness as described above. The first training method, which we call *gene-family guided splitting*, uses gene-family relationships to ensure that genes within the same family are not split between the training and testing sets. In this way, the model never sees a gene family in the testing set which it has already seen during the training process (Fig 1a). The second training method uses what we call *ortholog contrasts* (comparisons between pairs of orthologs) to eliminate evolutionary dependencies (Fig 1b). In addition to controlling for evolutionary relatedness, this method actually allows evolution to become an asset in the training process by leveraging whole genome duplication events and/or genetic differences between species, two things that would normally be a hindrance to such models. Using evolutionary relatedness is powerful because it allows one to understand and train of what has survived selection. Using deeper evolutionary divergence, between species rather than just within, allows for sampling thousands of years of mutagenesis and selective pressures.

**Figure 1.**
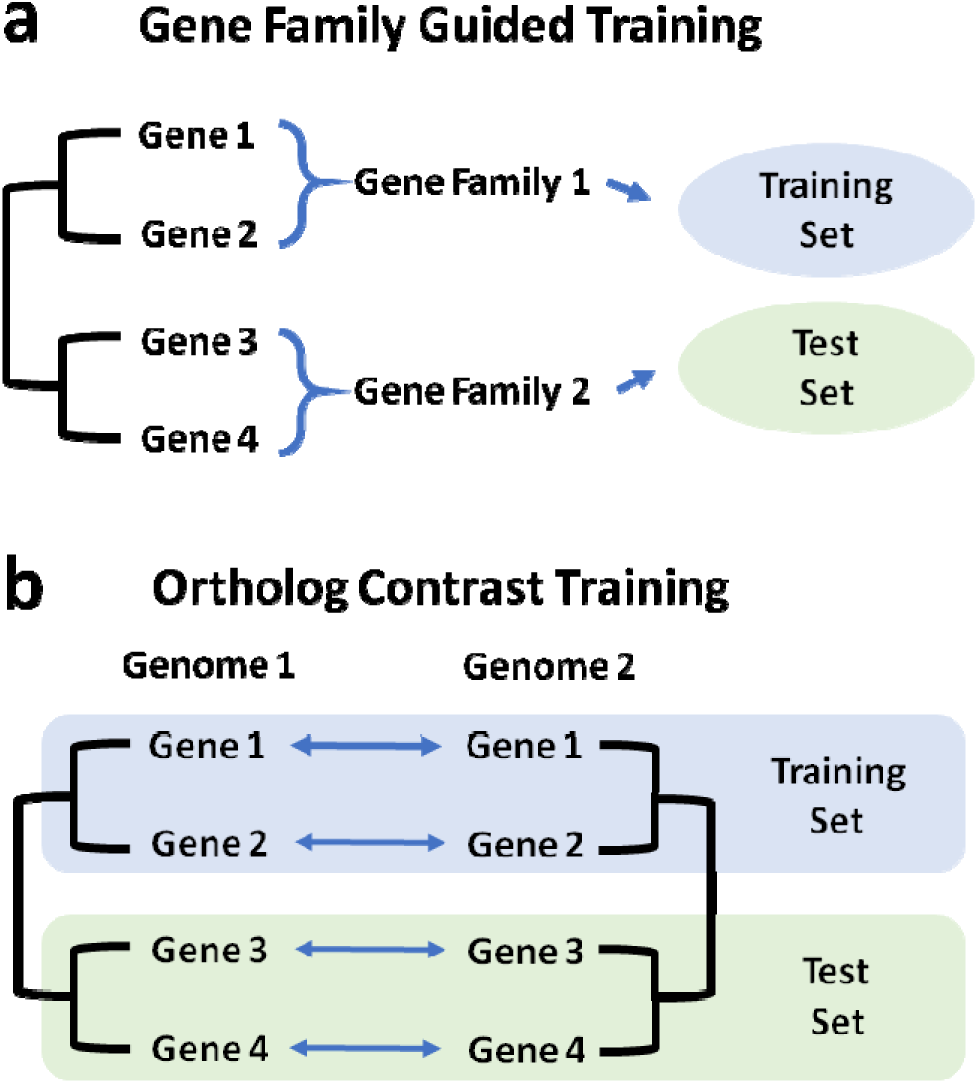
Evolutionarily informed strategies for deep learning. a) For prediction tasks involving a single species, genes are grouped into gene families before being further divided into training and test set, to prevent deep learning models from learning family-specific sequence features that are associated with target variables. b) For prediction tasks involving two species, orthologs are paired before being divided into training and test set, to eliminate evolutionary dependencies.

## RESULTS

### Differentiating between expressed and unexpressed genes based on DNA sequence

The first model was developed for the purpose of classifying genes as being expressed or not (zero or close to zero expression level). This model has been named the pseudo-gene model because of its ability to predict genes that are potentially pseudogenized and therefore lack expression. The pseudo-gene model also serves as a simple use case for the gene-family guided splitting approach. This model consists of a neural network composed of 6 convolutional layers and 3 fully-connected layers. It takes as input promoter and/or terminator sequences as defined in the online methods section and illustrated in Fig 2a. To generate the output of the model (i.e. a binary value representing whether a gene is expressed or unexpressed), a comprehensive atlas of gene expression in maize covering major tissues at various developmental stages was generated by applying a unified pipeline on 422 tissues from 7 RNA-Seq studies ^6–12^ (for full details, see the Online Methods, Supplementary Table 1, and Supplementary Table 2). The distribution of the maximum log-transformed Transcripts Per Million (logTPM) revealed a peak at the lower tail comprising unexpressed genes (4,562 genes with maximum logTPM<=1) along with normally distributed expressed genes (34,907 genes with maximum logTPM>1)(Fig. 1b).

**Figure 2.**
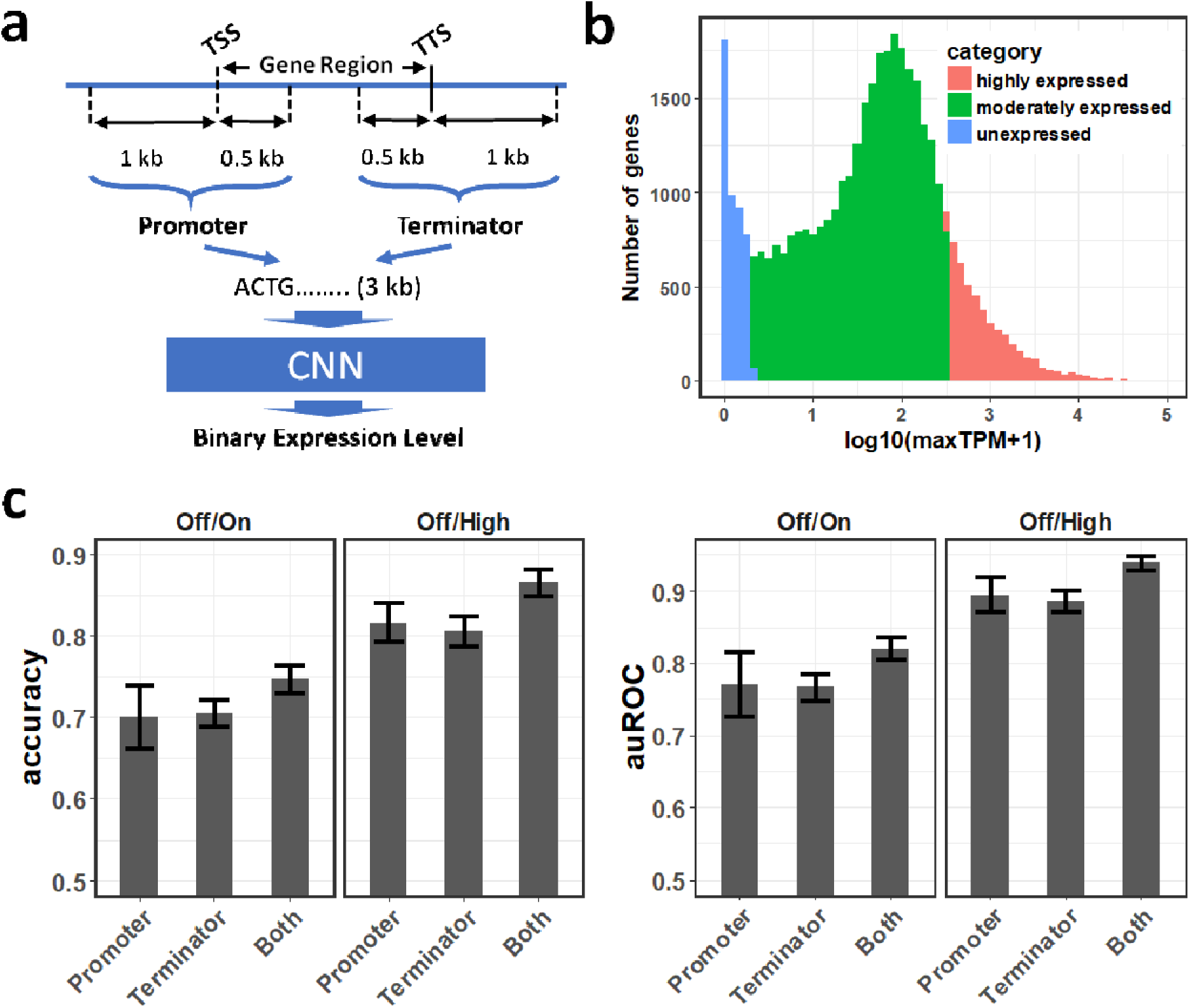
The architecture and performance of the pseudogene model. a) A schematic representation of the architecture of the pseudogene model. The model takes promoter and/or terminator sequences as the predictor to predict binary expression levels. b) A unified RNA-Seq data analysis pipeline was applied on 422 samples from 7 references ^6–12^ representing a comprehensive collection of maize tissues at diverse developmental stages. The log transformed maximal transcripts per million (TPM) over all samples was calculated for each gene, and used to represent the strength of predictor sequence. Shown is the distribution of log transformed maximal TPMs for all maize genes. Genes are categorized into unexpressed genes (blue), moderately expressed genes (green), and highly expressed genes (red). c) The accuracy and auROC of the pseudogene model trained on the Off/On gene set or the Off/High gene set, using promoters, terminators, or both promoter and terminator sequences as predictors. Error bars represent mean±sd from a gene family-guided 10 times 5-fold cross-validation procedure.

The number of expressed genes (34,907) and unexpressed genes (4,562) were highly imbalanced. Two approaches were used to handle the imbalance. First, expressed genes were divided into 4,562 highly-expressed (max TPM >= 342.9) genes and 25,783 intermediately-expressed genes (1 < max TPM < 342.9) and the model was trained to distinguish the 4,562 unexpressed genes from the 4,562 highly-expressed genes (off/high in Fig. 2c). As paralogous genes derived from more recent gene duplication events often share highly similar promoters or terminators, overfitting may potentially occur when highly similar paralogs are separated into training and testing sets. Moreover, as paralogs are often similar in their expression levels, separation of highly similar paralogs may force neural networks to learn gene family-specific sequence features, rather than sequence features that determine expression levels per se. To solve these problems, maize genes were divided into gene families (Online methods). The pseudo-gene model was trained on randomly selected families and tested on the remaining families not present in the training set (Supplementary Table 1).

The performance of the pseudo-gene model was evaluated using a 10 times 5-fold cross-validation procedure, and achieved an average predictive accuracy of 86.6% (auROC=0.94) when promoters and terminators were both used as the predictor. The average accuracy of the model reached 81.6% (auROC=0.89) and 80.6% (auROC=0.89) for promoters and terminators, respectively (Fig. 2c). Secondly, expressed genes were down-sampled to make them balanced with unexpressed genes. By using this approach, the model achieved an average predictive accuracy of 74.8% (auROC=0.82), 70.1% (auROC=0.77), and 70.6% (auROC=0.77) for both promoters and terminators, promoters only, and terminators only, respectively (off/on in Fig. 2c). The significance of the predictions was confirmed by the fact that shuffling predictor sequences while maintaining dinucleotide or single-nucleotide frequencies completely abolished the predictive accuracy of the models (Supplementary Table 2).

### Predicting which of two genes is more highly expressed using ortholog contrasts

The ortholog contrast model was designed to follow a simple approach derived from phylogenetics, where the most recent common ancestor of two closely related genes can be represented as a contrast between the two. Contrasting genes in this manner directly accounts for statistical dependencies between the genes that would otherwise hamper comparison with other genes ^13^. Building on this idea, the ortholog contrast method was here developed for comparing two genes from different genomes (or alleles from the same species) to each other and predicting the difference between the expression levels of the two (Fig 3a). When each gene is compared directly to its ortholog one can then compare that contrast value to the contrast values from other ortholog pairs without evolutionary dependence between them, hence enabling training and testing sets that are evolutionarily independent (Fig 1b, 3a).

**Figure 3.**
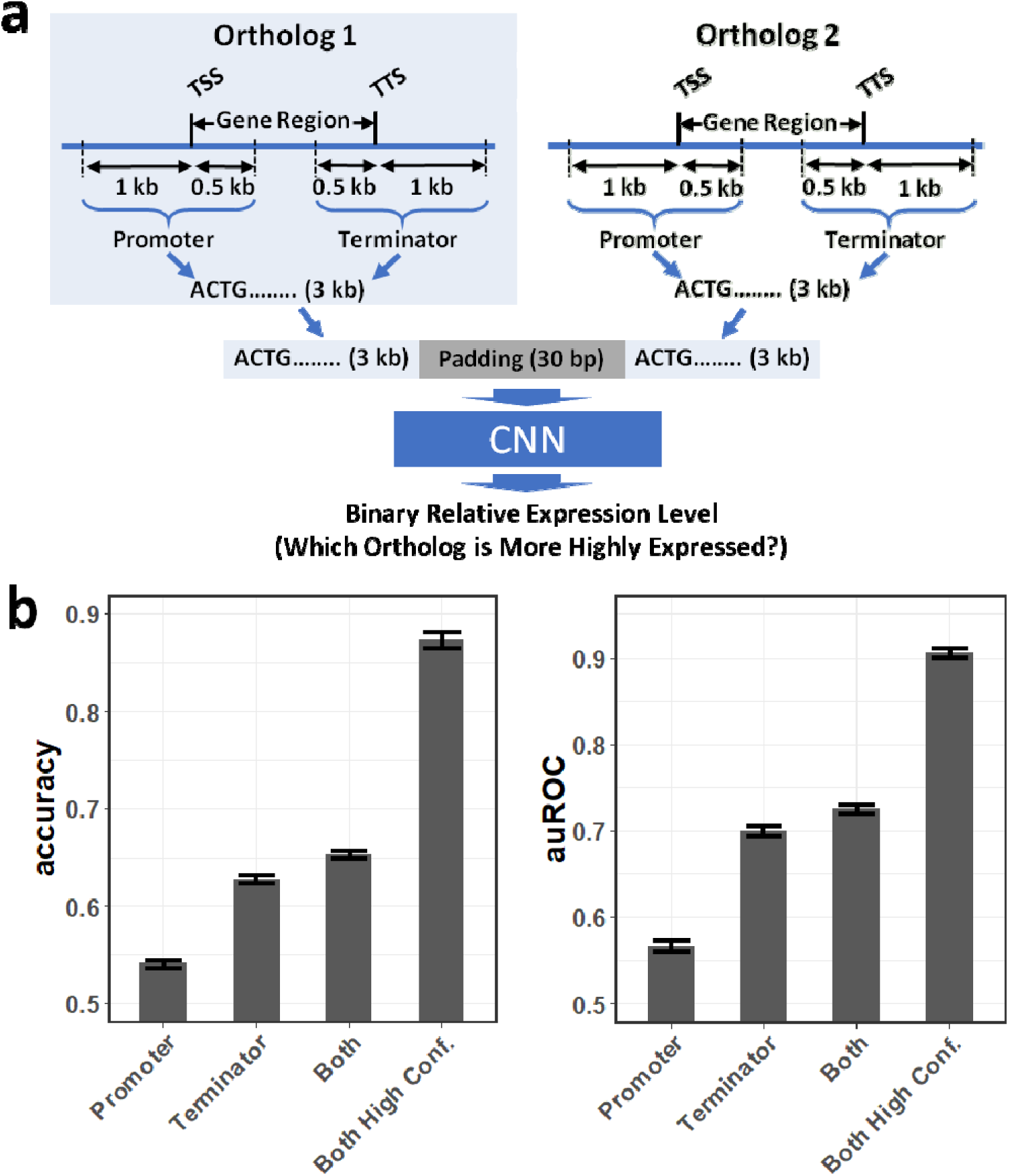
The architecture and performance of the ortholog contrast model. a) A schematic representation of the architecture of the contrast model. The model takes promoter and/or terminator sequences from two orthologous gene as the predictor and predicts the binary difference in expression level between the two. b) The accuracy and auROC of the ortholog contrast model trained using promoters, terminators, or both promoter and terminator sequences as predictors. Error bars represent mean±sd from 10 times 5-fold cross-validation. The bar “Both High Conf.” bar represents the models performance when genes for which the model has less than 0.5 confidence are dropped.

To further simplify prediction with the contrast model, we converted the values (the difference between the transcript abundance levels of the two compared genes) into simple binary form: zero if the first gene was more highly expressed than the second, and one in the opposite case. Orthologs with no expression difference between them were excluded. This simplification results in a model where the CNN is asked to determine which of two orthologs is most highly expressed. In reality, this question of deciding between two genes or alleles is actually what is most needed in applications like plant breeding and medicine.

The architecture used for this model was essentially the same as that used in the pseudo-gene model. The contrast model, when applied to comparisons between single-copy *Sorghum bicolor* and *Zea mays* orthologs (see online materials and methods), was able to predict never before seen pairs with an auROC value of 0.72 without filtering, and 0.91 with a test set that was filtered to only include samples for which the model had greater than 50% confidence in it prediction (Fig 3b). When applied to contrasts between the two *Z. mays* sub-genomes the model performed with auROC values of 0.63 and 0.71 without and with filtering respectively.

### Interpretation of CNN models reveals elements and motifs important for transcript abundance

Transcript abundance is concertedly determined by its production and degradation. Our current models cannot discriminate between these causes but could potentially do so in the future by training on GRO-seq (Global Run-On followed by high-throughput sequencing of RNA) ^14^, PRO-seq (Precision Run-On followed by sequencing) ^15^, or transcriptome-wide mRNA half-life data ^16,17^.

It has long been established that genomic sequences flanking coding sequences harbor important cis-elements that determine the transcription rate and/or the stability of transcripts ^18,19^. To identify these motifs/putative cis-elements, saliency maps ^20^, a way of determining which parts of the input sequence were most important to prediction, were applied on each of the above described models (Figs 4, 5). Interestingly, the different training methods (gene family-guided vs orthologs contrasts) resulted in very different but potentially complementary results as to which regions of the sequence were most important for the prediction task.

**Figure 4.**
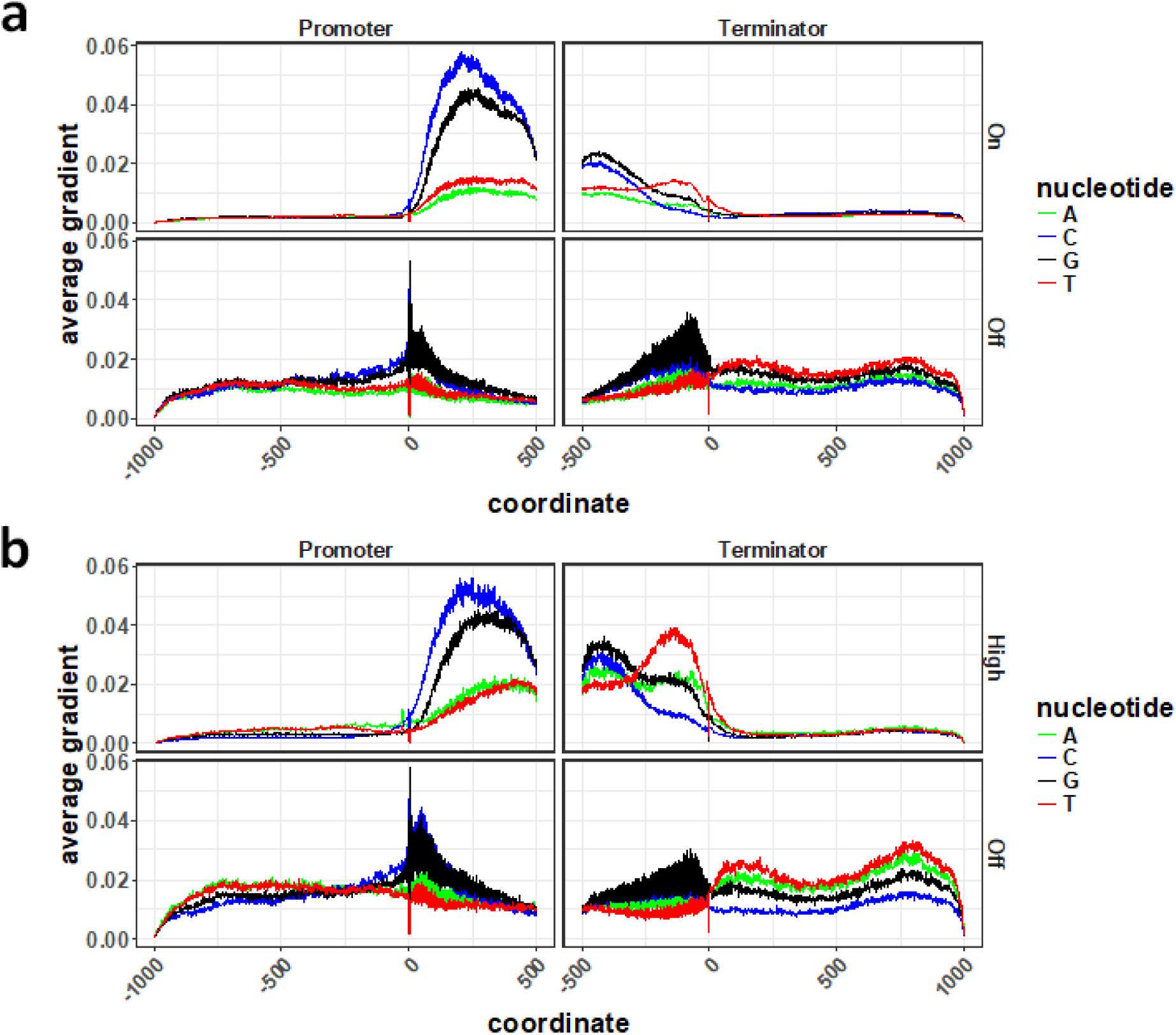
Averaged saliency map of the pseudogene model. Saliency map was calculated for the pseudo-gene model trained on either Off/On gene set (a) or the Off/High gene set (b). Saliency was averaged over non-pseudogenes (upper panels) and pseudogenes (lower panels), respectively. Only genes with correctly predicted expression levels were used for the calculation of saliency maps.

**Figure 5.**
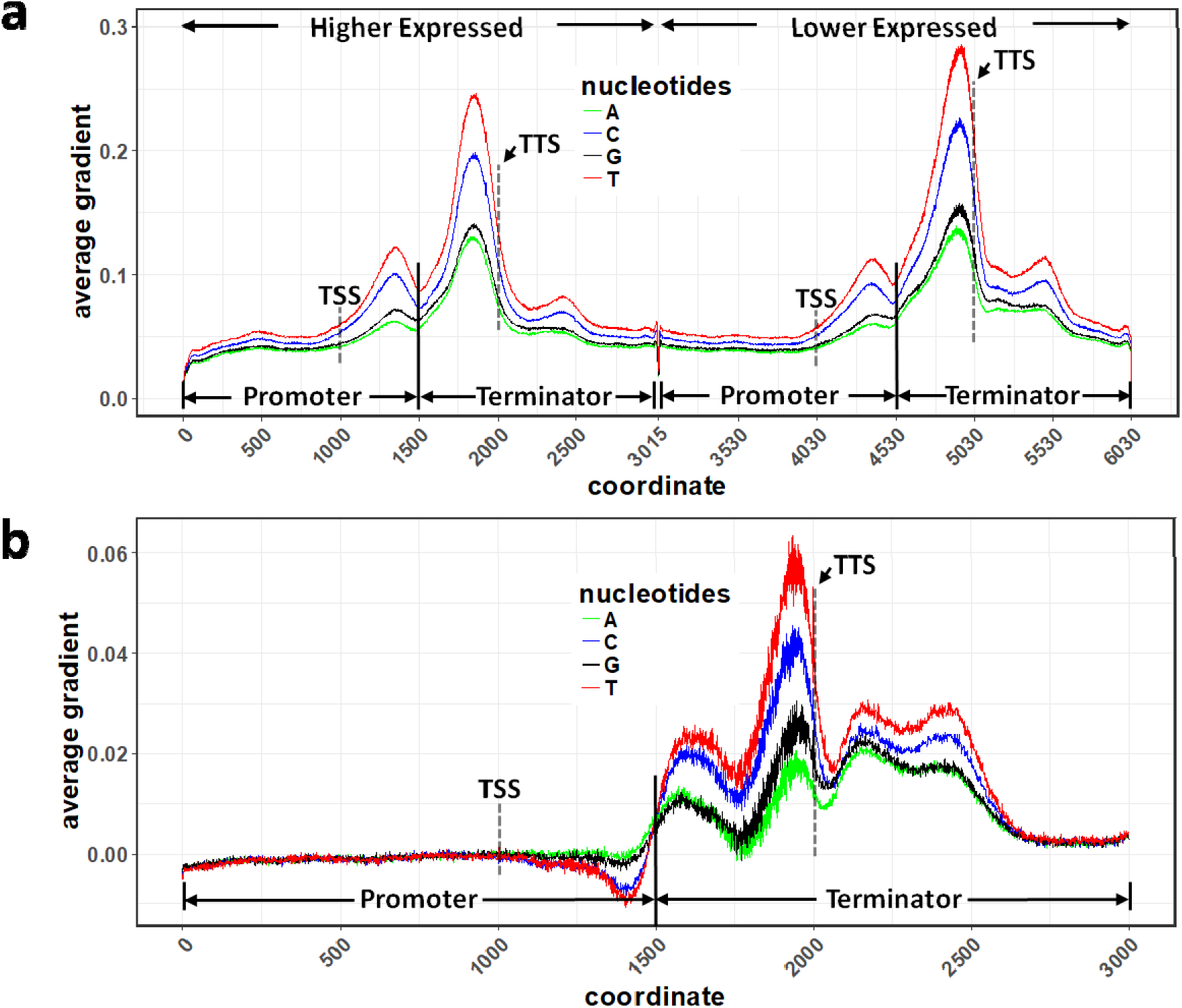
Averaged saliency maps of the ortholog contrast model. Saliency maps were calculated for the ortholog contrast model. a) True positive with ortholog 1 more highly expressed than ortholog 2. b) True positive lower expressed orthologs minus true positive higher expressed orthologs.

In all tested models, the 5’ and 3’ UTR regions of what we call promoter and terminator sequences were much more important than the regions just outside of the gene models (Figs 4, 5). Pseudo-gene models trained on the off/on data set (Fig 4a) and off/high data set (Fig 4b) resulted in similar saliency maps. In both cases, non-pseudogenes showed stronger signals in the promoter regions than in the terminator regions. This is in agreement with the accuracy values for the models shown above (Fig 2) where predictions based on the promoter sequences alone outperform those based on the terminator sequences alone. These results make sense based on what is currently known about cis regions that are important to gene expression.

For models trained using the ortholog contrast method, the results are somewhat different. In this case, the most heavily weighted areas of the input sequences were found in the terminator region (Fig 5a). This is consistent whether the gene in the first position is more highly expressed than the gene in the second position or visa versa. The greater importance of terminator regions is further shown by the fact that models run with only the terminator sequence perform better than those with only promoter sequences (Fig 3). The differences between expression values of the compared genes in the contrast model ranged from Log10 transformed TPM values of 0.12 to 2.80 with and average of 0.66. Log10 transformed TPM values for the off/high gene set in the pseudo-gene model ranged from 0 to 0.301 (with an average of 0.101) for the ‘off’ gene set, and ranged from 2.536 to 4.925 (with an average of 2.959) for the ‘high’ gene set.

While pseudo-genes (not expressed) in the family-guided approach showed very little signal in the promoter and terminator regions, the lower expressed genes in the contrast model actually showed very strong signals in these regions. In the case of the terminator region, lower expressed genes show higher values than highly expressed genes (Fig 5b), while the promoter regions show the opposite trend with higher saliency map values in the more highly expressed genes.

## DISCUSSION

### 3’ UTR potentially more important for small-scale changes in RNA abundance

There are a number of factors that might explain the differences in promoter and terminator importance between the the pseudo-gene and ortholog contrast models. First, the method of controlling for evolutionary relatedness differs between the models. The grouping of genes into families for the gene-family guided method relies on sequence similarity scores with subjectively defined cutoffs. This method is therefore potentially over or under controlling (or both in the case of different genes) for evolutionary relationships. The ortholog contrast model on the other hand fully controls for these relationships. Second, the pseudo-gene model is restricted to a single species while the contrast was applied both within species (though between sub-genomes) and across species. Lastly, and perhaps most likely, is that the two models are focused on different categories of gene expression and genes which have experienced different types of evolutionary constraint. The pseudo-gene model includes a bimodal distribution of genes that are expressed (highly or moderately) and genes that are not expressed, while the contrast model mostly contains genes that are expressed as some level (it likely does not include many pseudo-genes). Given that small mutations in the promoter region could conceivably inhibit enzyme binding and be responsible for drastic changes in gene expression (such as turning expression on or off), perhaps these types of mutations are responsible for the high predictive importance of promoter regions in the pseudo-gene model. The contrast model on the other hand is limited to genes with matching syntenic orthologs between distinct genomes. These genes will be highly conserved, and under strong purifying selection, meaning that large scale expression changes are unlikely to be tolerated. Small-scale, fine-tuning adjustments of expression level on the other hand are more likely to be present in these highly conserved genes. We therefore hypothesize that the terminator region (particularly the 3’UTR) plays a more important role in small-scale, fine-tuning of RNA abundance levels than does the promoter (particularly the 5’ UTR) region. The 3’ UTR’s importance in RNA stability suggests one possible mechanism for this tuning.

### Strengths and weaknesses of different models and training approaches

We have demonstrated the validity of two different approaches for mRNA expression prediction. Both approaches incorporate methods for dealing with evolutionary relatedness between genes within a predictive framework. Each approach has strengths and weaknesses, and which approach one choses will depend on the datasets available and the types of predictions/ biological insights desired. For datasets or questions that are limited to a single sample from a single diploid species the gene-family guided approach is the most suitable option, because orthologs contrasts are not possible. This method may also be most appropriate in situations where one wants to specifically identify gene that are unlikely to be expressed. The gene-family guided method also has the benefit of being somewhat simpler to understand and interpret, however, one must define gene-families, and that process is subjective and sensitive to parameter choices. In situations where multiple species or genotypes are involved, the contrast method is likely to be the method of choice. The contrast method is also the better choice when one wishes to compare orthologs/alleles that are both expressed at a moderate to high level.

### Conclusions and future applications

In the current study, CNN models have been successfully applied to the prediction of mRNA abundance under several distinct scenarios. The models are able to predict a gene’s on/off state as well as which of two compared orthologs is more highly expressed. These models feature two very different approaches for handling evolutionary relationships between genes, gene-family guided splitting and ortholog contrasts. While not demonstrated here, the contrast model should in theory be extendable to determining which of two alleles in a population is more highly expressed, an application with clear utility in breeding and medicine.

In the future, it is hoped that gene-family guided splitting, ortholog contrasts, and other potential strategies will be applied to deep learning models in various areas of biology. The potential of deep learning to increase our understanding of and prediction within biological systems is enormous. Further creative strategies for focusing these methods on biologically relevant information and controlling for confounding biological factors will be critical the success of deep learning in biology.

## METHODS

### DNA sequence encoding and RNA-Seq data collection and processing

The *Zea mays* B73 reference genome ^21,22^ was downloaded from Ensembl Plant Release 31 (http://plants.ensembl.org). The newest B73 reference genome, AGPv4 ^23^, was not used in this study due to known issues with the the 3’ UTR annotations. Version 3.1.1 of the *Sorghum bicolor* genome ^24^ was downloaded from Phytozome (https://phytozome.jgi.doe.gov). The TSS and TTS sites are not explicitly annotated in these genomes, so the TSS was taken to be the start coordinate of the gene and the TTS the end coordinate of the gene. DNA sequences were transformed into one-hot form using custom scripts (see bitbucket repository).

To train models predicting unexpressed genes, 452 samples from 7 references ^6–12^ representing a comprehensive collection of maize tissues at diverse developmental stages were used. The 452 RNA-Seq samples were downloaded from NCBI Sequence Read Archive (SRA). The downloaded sra files were converted to fastq format using fastq-dump in SRA Toolkit (version 2.8.2, https://github.com/ncbi/sra-tools). Reads were quality trimmed by sickle (version 1.33, https://github.com/najoshi/sickle), followed by quality checking with FastQC (version 0.11.5, https://www.bioinformatics.babraham.ac.uk/projects/fastqc/). The clean reads were then aligned to the maize genome by HISAT2 (version 2.1.0)^25^. Gene-level raw read counts were normalized using Transcript per million (TPM) by stringtie (version 1.3.3)^26^. Among the 452 samples, 26 samples with less than 5 million reads and 4 samples with less than 50% of total alignment rate were excluded from downstream analysis, leaving 422 samples. Samples are summarised in Supplementary Table 1, gene expression levels are listed in Supplementary Table 2.

Some gene models in the V3 annotation do not contain 5’ or 3’ UTRs, leaving discernible start and stop codons at TSS and TTS. To circumvent the model learning from these simple sequence features, the first three nucleotides downstream of the TSS and the last three nucleotides upstream of the TTS were masked. In both off/high and off/on comparisons, using longer promoters (from −2500 bp to +500 bp with respect to the TSS) or terminators (from −500 bp to +2500 bp with respect to TTS) did not improve predictive accuracy.

The *Z. mays* samples used for testing the ortholog contrast model come from the B73 Shoot data published by Kremling et al. 2018 ^27^. The corresponding *S. bicolor* data was generated and processed as described in that paper and can be found on NCBI SRA under accession number XXX. Both Z. mays and S. bicolor data were additionally processed together using the DESeq2 fragments per million normalization. All scripts associated with the analyses were deposited on BitBucket (XXX)

### Categorizing maize genes into gene families and family-guided splitting of training and testing data sets

*Z. mays* genes were divided into gene families using a previously described pipeline ^28^ with modifications. An all-by-all BLAST was conducted on maize proteome sequences to evaluate pairwise similarity between maize proteins. As one gene may encode multiple protein isoforms, the result of the blast search was collapsed to gene-level similarity by an in-house R script. Then, an in-house python script was used to build a graph with nodes representing genes and edges connecting paralogous genes. This graph was further divided into clusters (i.e. gene families) by the Markov Clustering Algorithm (MCL) implemented in the markov_clustering package in python with default parameters except that inflation was set to 1.1. If a gene was not assigned to any gene family, it was considered as a family that contains only a single member. Each gene family was assigned an index (Supplementary Table 3). For family-guided 5-fold cross-validation, gene families were randomly partitioned into 5 sub-samples with equal numbers of families. In each iteration, 1 subsample is retained as the test data, while the remaining 4 sub-samples were used as training data (Supplementary Table 3).

### Syntenic ortholog contrasts

Syntenic orthologs between the *Z. mays* and *S. bicolo*r reference genomes were obtained from a previous publication ^11^. The training, validation, and testing sets were also divided by gene family in the same way as described for the pseudo-gene model above. In order to feed two genes at a time to the model, the gene sequences were first converted into one-hot form. Each base-pair, and a missing base-pair character, were included in the encoding. A column of all zeros was used as an additional class specific to padding characters. The two ortholog sequences being compared were then concatenated together with a block of all zero columns equivalent to 30 base-pairs in between them. Different lengths of padding between the two sequences were tried, but 30 base-pairs worked well and was used for all reported analyses. Transcript abundance values were log2 scaled and then normalized by percentage rank. In cases where multiple transcripts had the same expression values, the average rank value was assigned to all.

## ACKNOWLEDGMENTS

Thank you to James Schnable for the maize and sorghum ortholog lists. This material is based upon work supported by the NSF Postdoctoral Research Fellowship in Biology under Grant No. 1710618 (JDW), the NSF Plant Genome Research Program under Grant No. 1238014 (ESB), and the Tang Cornell-China Scholars Program (HW). Additional support comes from the United States Department of Agriculture - Agricultural Research Service.

## AUTHOR CONTRIBUTIONS

JDW, ESB, MKMG, GR and KAK conceived of and implemented the ortholog contrast method. HW and ESB conceived of, implemented, and refined the pseudogene model. RV and KAK generated and processed 3’ RNA expression data. HW reanalyzed maize RNA-Seq data from NCBI Sequence Read Archive.

